# Onset of natural selection in auto-catalytic heteropolymers

**DOI:** 10.1101/204826

**Authors:** Alexei V. Tkachenko, Sergei Maslov

## Abstract

Reduction of information entropy along with ever-increasing complexity are among the key signatures of living matter. Understanding the onset of such behavior in early prebiotic world is essential for solving the problem of origins of life. To elucidate this transition, we study a theoretical model of information-storing heteropolymers capable of template-assisted ligation and subjected to cyclic non-equilibrium driving forces. We discover that this simple physical system undergoes a spontaneous reduction of the information entropy due to the competition of chains for constituent monomers. This natural-selection-like process ultimately results in the survival of a limited subset of polymer sequences. Importantly, the number of surviving sequences remains exponentially large, thus opening up the possibility of further increase in complexity due to Darwinian evolution. We also propose potential experimental implementations of our model using either biopolymers or artificial nano-structures.

## Introduction

The second law of thermodynamics states that the entropy of a closed system increases with time. Life represents a remarkable example of the opposite trend taking place in an open, non-equilibrium system [1]. Indeed, both information and thermodynamic entropies decrease in the course of Darwinian evolution reflecting ever-increasing complexity of living organisms and their communities [2]. Interestingly, both the second law and the concept of entropy were introduced by Clausius in 1850s, roughly at the same time as Darwin developed and published his seminal work. A century later the connection between life and entropy was highlighted in the classical work of Schrödinger titled “What is life?” [3]. According to him, living systems are characterized by their ability to “feed on” and store the negative entropy (which he referred to as “negentropy”) [2]. In the same work, he effectively predicted the existence of information-storing molecules such as DNA. Soon after, Brillouin established [4] the connection between the thermodynamic negentropy and its information cousin defined by Shannon [5].

The emergence of life from non-living matter is one of the greatest mysteries of fundamental science In addition, the search for artificial self-replicating nano-and micro-scale systems is an exciting field with potential engineering applications [6, 7, 8, 9]. The central challenge in both of these fields is to come up with a simple physically-realizable self-replicating system obeying the laws of thermodynamics, yet ultimately capable of Darwinian evolution when exposed to non-equilibrium driving forces.

Chemical networks of molecules engaged in the mutual catalysis have long been considered a plausible form of prebiotic world [10, 11, 12, 13]. Furthermore, a set of mutually-catalyzing RNA-based enzymes (ribozymes) is one of the best known examples of experimentally realized autonomous self-replication. This is viewed as a major evidence supporting the RNA-world hypothesis (see e.g. Refs. [14]-[18]). The ribozyme activity requires relatively long polymers made of hundreds of nucleotides with carefully designed sequences whose spontaneous emergence by pure chance is nearly impossible. Thus, to make the first steps towards explanation of the origin of life, one needs to come up with a much simpler system capable of spontaneous reduction of the information entropy, that would ultimately set the stage for Darwinian evolution e.g. towards functional ribozymes and/or autocatalytic metabolic cycles.

**Figure 1:**
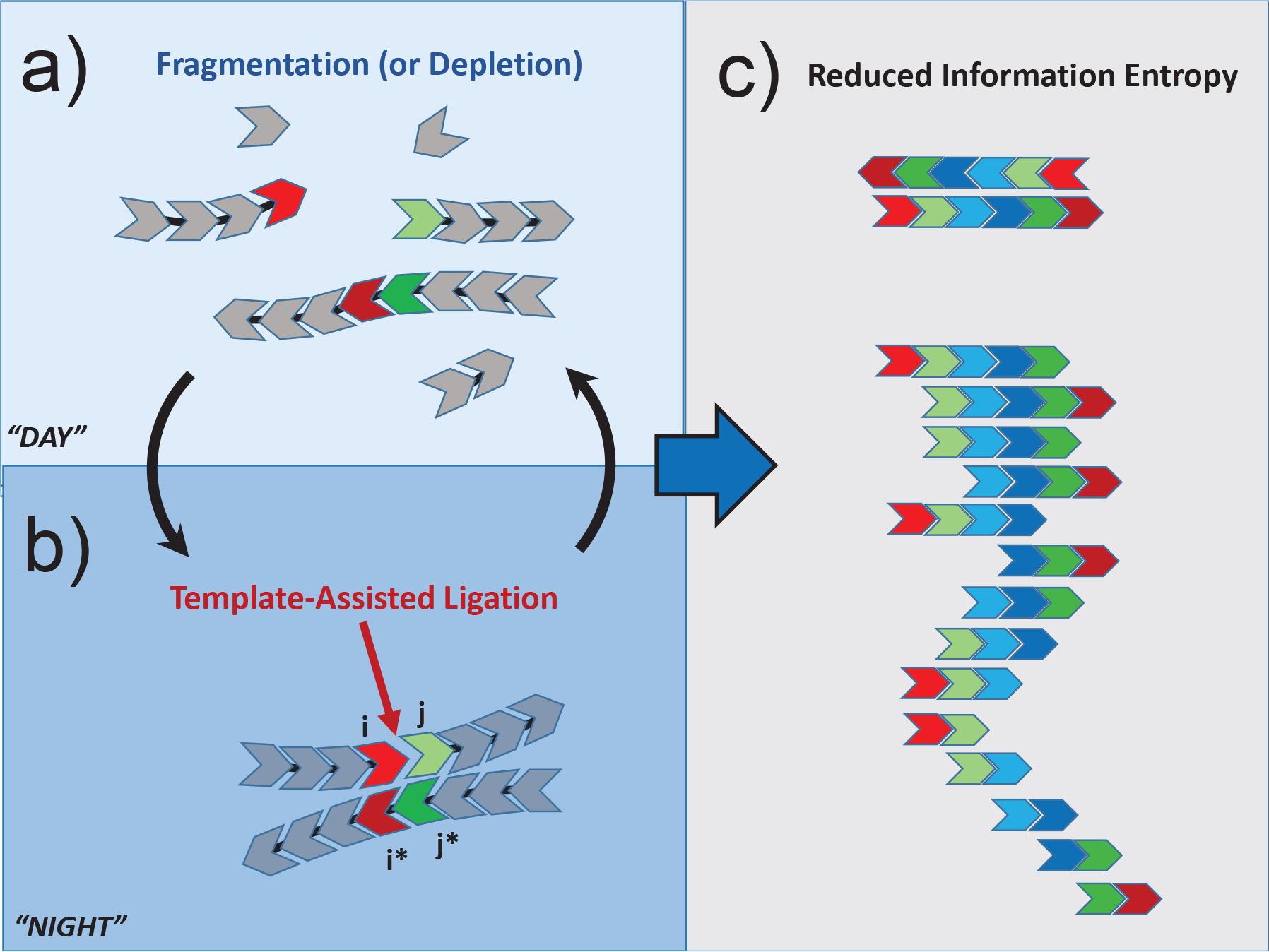
A conceptual representation of our model in which the system is cycled between day, a), and night, b), phases. During the night phase polymer chains undergo template-assisted ligation joining left and right ends *i* and *j* respectively to form a new 2-mer *ij*. This process is assisted by a complementary 2-mer *j*\**i*\*. The process results in a reduced information entropy of chain sequences dominated by a small subset of 2-mers illustrated in c).

A promising candidate for such mechanism is provided by template-assisted ligation. In this process pairs of polymers are brought together via hybridization with a complementary template chain and eventually ligated to form a longer chain (see Fig. 1a-b). Unlike the non-templated reversible step-growth polymerization used in Ref. [19], this mechanism naturally involves the transmission of an information from a template to the newly-ligated chain, thus opening an exciting possibility of long-term memory and evolvability. An early conceptual model involving template-assisted polymerization was proposed by P. W. Anderson and colleagues [20, 21]. It has also been a subject of several more recent experimental and theoretical studies [24, 25, 22]. In particular, the model by Hordijk et al. [22] makes a connection between the classical Kauffman model of autocatalytic sets [12] and polymer systems capable of template-assisted ligation (see also [23] for further development of that approach). Recently, we theoretically established [26] that a cyclically-driven system of this type is capable of producing long, mutually-catalyzing chains starting from a primordial soup dominated by monomers. In the current study we focus on statistics of sequences of these chains and discover that the dynamics of the system naturally results in a dramatic reduction of the information entropy in the sequence space.

## Model

Here we further develop the model introduced in Ref. [26]. It describes the emergence of heteropolymers out of the “primordial soup” of monomers by virtue of template-assisted ligation. Our system is driven out of equilibrium by cyclic changes in physical conditions such as temperature, salt concentration, pH, etc. (see Fig. 1).

We consider a general case of information-coding heteropolymers composed of *Z* types of monomers capable of making *Z*/2 mutually complementary pairs. Polymerization occurs during the “night” phase of each cycle when existing heteropolymers may serve as templates for ligation of pairs of chains to form longer ones. When the end groups of two substrate chains are positioned next to each other by the virtue of hybridization with the template, a new covalent bond connecting these end groups is formed at a certain rate (see Fig. 1b). During the “day” phase of each cycle all hybridized pairs dissociate, and individual chains are fully dispersed (see Fig. 1a).

One of the key results of our previous work [26] is the existence of the optimal hybridization overlap length *k*_0_ for template-substrate binding. In this work, for the sake of simplicity we assume that *a single pair* of complementary monomers is sufficient to bind a substrate to the template. This can be interpreted as if each of *Z* monomers in the present model is in fact a “word” composed of *k*_0_ smaller elementary letters, such as e.g. RNA or DNA bases. Within this interpretation, the number *Z* of such “composite monomers” can be exponentially large: *Z* = *z*^*k*0^, where *z* is the number of elementary letters (*z* = 4 in the case of RNA). As in [26] we ignore the process of spontaneous, non-templated ligation [25, 19].

In our model monomer types are labeled in such a way that type *i* is complementary to type *i*\*. One of the key concepts in our analysis is that of a “2-mer” *ij* referring to a monomer *i* immediately followed by the monomer *j* and *found anywhere within any heteropolymer*. Note that, similar to DNA/RNA complementary strands, polymers in our system are assumed to be directional and anti-parallel when hybridized to each other. Therefore, a 2-mer *j*\**i*\* formed from monomers *j*\* and *i*\* is complementary to the 2-mer *ij*. It can serve as a template catalyzing the ligation of two substrate chains with monomers *i* and *j* located at their appropriate ends. (see Fig. 1b).

Let *d*_*ij*_ denote the overall concentration of 2-mers of type *ij*, i.e. the total number of consecutive monomers of types *i* and *j* found anywhere within any chain, divided by the volume of the system. We will refer to the *Z* × *Z* matrix formed by all *d*_*ij*_ as the 2-mer matrix. Let *l*_*i*_ denote the concentration of all chains ending with a monomer of type *i*, while *r*_*j*_ - the concentration of all chains starting with a monomer of type *j*. When two ends *i* and *j* of such chains meet due to hybridization with a complementary template *j*\**i*\*, they are ligated at a certain rate to form a new 2-mer *ij*. We describe this process by a three-body mass-action term λ_*ij*_ · *l*_*i*_(*t*) · *r*_*j*_(*t*) · *d*_*j*\**i*\*_(*t*). Here λ_*ij*_ is the ligation rate averaged over the duration of the day-night cycle with the understanding that it happens only during the night phase. 2-mers *ij* in our system are assumed to spontaneously break up at a rate *β*_*ij*_. Thus we extend the original model by introducing an explicit sequence dependence of ligation (λ_*ij*_) and breakage (*β*_*ij*_) rates. Master equations, describing the slow dynamics in our system occurring over multiple day/night cycles, are:

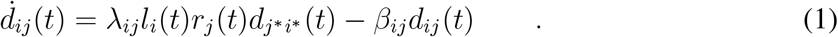

This mass-action description implies that our system stays well below the saturation regime during the night phase. In other words, we assume that template-substrate hybridization probability is determined by the association rate and not by the competition of multiple different substrates for the same binding site on a chain. This is realized when the duration of the night phase of the cycle is shorter than the typical association time for hybridization. Importantly, this regime also ensures that there is no template poisoning, i.e. the probability of two complementary 2-mers binding each other (and thus loosing their catalytic activity) remains low.

One could write a similar set of kinetic equations describing the dynamics of concentrations of “left” and “right” ends of chains, *l*_*i*_(*t*) and *r*_*i*_(*t*). Instead, we use the conservation of overall concentrations of monomers of each type to obtain the explicit algebraic expressions for *l*_*i*_(*t*) and *r*_*i*_(*t*) in terms of the 2-mer matrix:

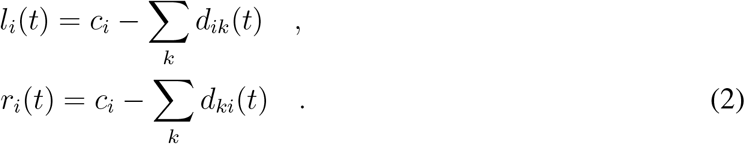

Here *c*_*i*_ is the overall concentration of monomers of type *i* in the pool, both free and bound. At the start only free monomers are present (*d*_*ij*_ = 0), and thus the initial conditions are given by *l*_*i*_(0) = *r*_*i*_(0) = *c*_*i*_.

Our model allows for an alternative interpretation that does not involve breakage of intrapolymer bonds. Specifically, as shown in the Supplementary Materials (SM), Eqs. (1-2) also describe a system subject to uniform dilution at rate *β* and the influx of fresh monomers *ϕ*_*i*_ = *β* · *c*_*i*_. In the light of this interpretation, below we focus on the case of all *β*_*ij*_ equal to each other. Without loss of generality, they can all be set to unity: *β*_*ij*_ = 1. This defines the fundamental timescale in our system as either the average lifetime of a single bond or the inverse of the dilution rate.

## Results

### Spontaneous entropy reduction

In our previous study [26] we worked within random sequence approximation. If all monomers have identical total concentrations *c*_*i*_ = *c*, this approximation corresponds to all matrix elements *d*_*ij*_(*t*) being equal to each other. For general initial conditions these elements would be proportional to *c*_*i*_ · *c*_*j*_. The key hypothesis proposed but not tested in Ref. [26] is that the system dynamics would eventually favor the survival of a subset of the “fittest” sequences at the expense of the others, thus breaking the random sequence approximation. Here we test this hypothesis by simulating the dynamics of the model given by Eqs. (1-2) with *Z* = 20. We start with a system characterized by a weak variation in individual ligation rates λ_*ij*_ and concentrations *c*_*i*_. We choose them from a log-normal distribution with their logarithms having standard deviation 0.1 and means 0 and log 3 respectively. Our choice of parameters is motivated by the need to understand the the limit of infinitesimally weak variation of rates and concentrations. For this combination of parameters, the Eqs. 1 are initially linearly unstable with respect to formation of all 2-mers. However, no 2-mer would be formed until either it or its complementary partner is present in the system at least in some infinitesimal “seed” concentration. Once such seed is introduced, the corresponding pair of mutually complementary 2-mers *ij* and *j*\**i*\* would be exponentially amplified. In our simulations we used the same small seed concentration of 10^−4^ for all *Z*^2^ 2-mers

The key parameter we use to quantify the emergent complexity in our system is the information entropy of 2-mers based on their relative concentrations 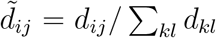 and defined in the standard Boltzmann-Shannon manner:

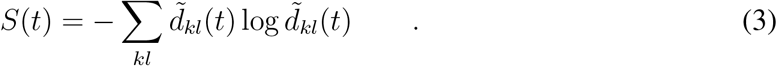

Fig. 2 shows the time dependence of this entropy in 5 different realizations of λ_*ij*_ and *c*_*i*_. The entropy starts at its maximal value log(*Z*^2^), and after a brief dip followed by a rebound, it steadily *declines* as a function of time. Such behavior is a remarkable manifestation of the non-equilibrium nature of our system, as the entropy changes in the direction opposite to that dictated by the second law of thermodynamics. To reveal the source of this entropy dynamics, in Figs. 2b and 2c we show the 2-mer matrix at two time points during our simulations. At *t* = 2 all of the 2-mers have grown up from their seed to substantial concentrations. Remarkably, the subsequent dynamics leads to a *complete extinction* of the majority of 2-mers ultimately giving rise to the 2-mer matrix at *t* = 8000 shown in Fig. 2c. The time dependence of the logarithm of the number of surviving 2-mers is shown as red lines in Fig. 2a. The ultimate number of survivors 36 ± 4 is just below 2*Z* = 40 (out of *Z*^2^ = 400) represented by the lower horizontal dotted line at log 2*Z* in Fig. 2a.

### Competition between 2-mers and the number of survivors

The observed behavior can be understood from the analysis of Eqs. (1-2). For a fixed set of concentrations *l*_*i*_ and *r*_*i*_, Eqs. 1 form a set of linear kinetic equations with respect to 2-mer concentrations *d*_*ij*_. Furthermore, this set of *Z*^2^ equations breaks into independent blocks of equations describing the dynamics of mutually complementary 2-mers *d*_*ij*_ and *d*_*j*\**i*\*_. For a small subset of self-complementary 2-mers *ii*\*, occupying a diagonal of the 2-mer matrix, such a block is represented by a single equation. In all other cases it involves a pair of equations for *d*_*ij*_ and *d*_*j*\**i*\*_ coupled via a 2×2 matrix:

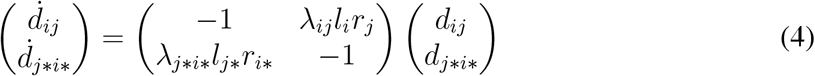

**Figure 2:**
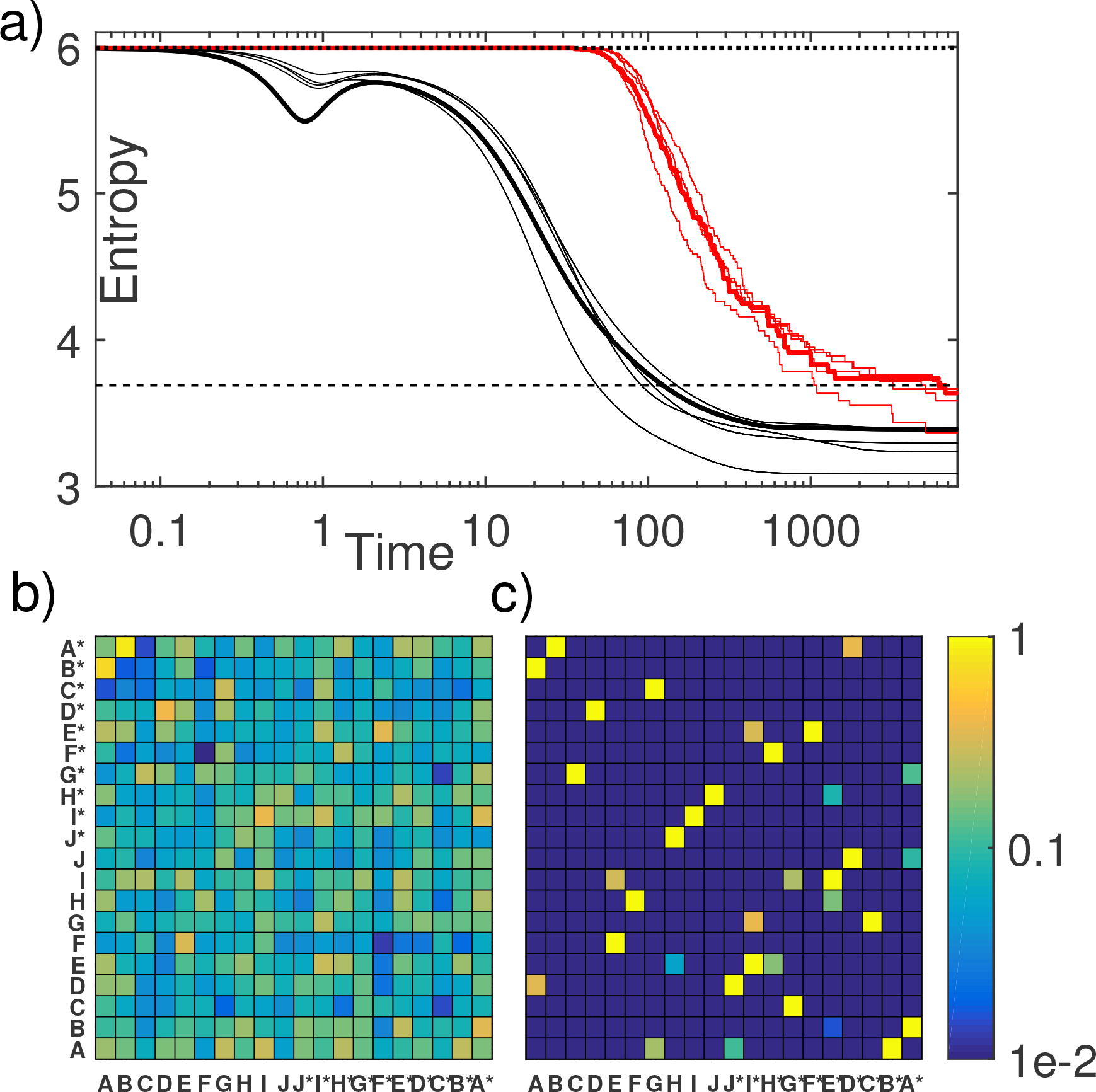
a) The information entropy *S* (black lines) given by Eq. 3 and the natural logarithm of the number of surviving 2-mers *N* (red lines) plotted vs time in 5 different realizations of our model with logarithms of both λ_*ij*_ and *c*_*i*_ normally distributed with standard deviation of 0.1 and means of log 1 and log 3 respectively. b) The heatmap visualizing log_10_ of concentrations of 2-mers at *t* = 2 (the second maximum of the entropy) in one of these realizations highlighted by thick black and red lines in panel a). c) The same heatmap in the steady state at *t* = 8000 where the entropy is saturated at its lowest point.

Because the trace of the matrix is always negative, at least one of the eigenvalues has to have a negative real part, while the real part of the other one could be positive, negative, or zero depending on the value of the matrix determinant Δ_*ij*_. A negative value of the determinant Δ_*ij*_ < 0 corresponds to a positive eigenvalue and hence to the exponential growth of two complementary 2-mer concentrations observed at the initial stage. As growing 2-mers gradually deplete *l*_*i*_ and *r*_*j*_, Δ_*ij*_ increases and may eventually turn positive. In this case both eigenvalues become negative. This triggers the exponential decay of concentrations and ultimate extinction of the corresponding pair of 2-mers. A small subset of 2-mers survive and reach the steady state. For these survivors, the determinant *has to become exactly zero*: Δ_*ij*_ = 0. These conditions for surviving 2-mers can be rewritten as

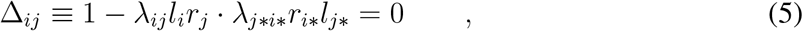

while for all extinct 2-mers Δ_*ij*_ > 0.

Now we can put the upper bound on the number of surviving 2-mers in the steady state of the system. This is accomplished by comparing the total number of constraints given by Eq. 5 to the number of independent variables. Since matrix 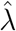 is fixed, the only variable parameters in Eqs. 5 are the left and right end concentrations *l*_*i*_(*t*) and *r*_*i*_(*t*). While naively, the number of such variables is 2*Z*, Eqs. 5 always contain them in combinations *l*_*i*_ · *r*_*i*\*_. Therefore, for the purpose of our counting argument, only these *Z* products should be considered as independent variables. The number of constrains Eqs. 5) that are simultaneously satisfied cannot be greater than *Z*. Each of these equations corresponds to either a pair of mutually complementary 2-mers or to a single self-complementary 2-mer. Denoting the total number of surviving 2-mers as *N*, and the number of self-complementary surviving 2-mers as *N*_*sc*_, the number of equations for surviving 2-mers is given by (*N* + *N*_*sc*_)/2. Thus the upper bound on the number of surviving 2-mers is given by

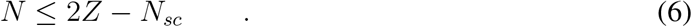

Note that for large *Z* the number of surviving 2-mers is dramatically lower than *Z*^2^ - the total number of possible ones. This explains the entropy reduction observed numerically (see Fig. 1). The parameters of the system were chosen in such a way that initially all *Z*^2^ 2-mers grow exponentially. Since the rate of this early exponential growth depends on λ_*ij*_ and *c*_*i*_, it differs from one 2-mer to another. This results in a transient behavior where the inhomogeneity of 2-mer concentrations is amplified giving rise to an early decrease in entropy (see the dip around *t* = 1 in Fig. 2a). As concentrations *l*_*i*_ and *r*_*i*_ start to get gradually depleted, the growth saturates, giving time for slower-growing 2-mers to catch up with the faster-growing ones around *t* = 2 (see Fig. 2b). As a consequence, the entropy recovers close to its maximal value. After that, a new process starts in which 2-mers actively compete with each other for remaining left and right ends. When the determinant Δ_*ij*_ for a particular pair of 2-mers *ij* and *j* * *i*\* changes its sign to positive, that pair of 2-mers starts to exponentially decay and eventually goes extinct. This process continues until the number of remaining 2-mers *N* (for which Δ_*ij*_ = 0), falls below the upper bound given by the Eqs. 6. These surviving 2-mers are visible in the heatmap in Fig. 2C.

### Graph-theoretical representations

A useful visualization of the emergent state of the system is a so-called de Bruijn graph shown in Fig. 3a. It represents each of *Z* monomer types as a vertex, and each of *N* surviving 2-mers *ij* as a directed edge connecting vertices *i* and *j*. The weight of every edge is proportional to the steady state 2-mer concentration *d*_*ij*_. A de Bruijn graph is a common representation of heteropolymer ensembles such as e.g. DNA sequences of all the chromosomes in an organism. It is straightforward to construct it from a known pool of sequences. However, the inverse problem of reconstruction of the statistics of a sequence pool from the de Bruijn graph is highly non trivial. Each of the polymers in the pool can be represented as a walk on this graph. The simplest case is when the consecutive steps of this walk are uncorrelated with each other. This means that the walk is a random Markov process with the probability of a step *i* → *j* given by *d*_*ij*_/*c*_*i*_, while the probability of termination of a polymer at the vertex *i* given by 1 − Σ_*j*_ *d*_*ij*_/*c*_*i*_ = *l*_*i*_/*c*_*i*_. Note that the entropy defined above and plotted in Fig. 2a is *exactly* the information entropy of a pool of polymer sequences generated by such Markov process [28]. Longer-range correlations are not captured by the present model, but can in principle could emerge due to the effects outlined in the Discussion section, leading to a further reduction of the information entropy in the system.

The de Bruijn graph can be complemented by another, more compact, graphical representation specific to our system. Since mutually complementary 2-mers appear in pairs *ij* and *j*\**i*\* (with the exception of self-complementary 2-mers *ii*\*), each such pair can be depicted as a single undirected edge connecting vertices *i* to *j*\*. In this representation each edge represents two 2-mers, while each vertex *i* stands for either *i* or *i*\* monomer, depending on whether it is the first or the second letter within the 2-mer. For our system this undirected graph (see 2b) has a number of remarkable properties derived in SM. First, it is a so-called “pseudoforest” [27]: each of its individual connected components contains no more than one cycle. This allows us to refine and give a topological interpretation to Eqs. 6: *N* = 2*Z* − *N*_*sc*_ − 2*N*_*trees*_, where *N*_*trees*_ is the number of trees (components without cycles) in the pseudoforest. Second, only the odd-length cycles (1,3,5, etc.) are allowed in this graph.

Fig 3c shows the distribution of the number of surviving 2-mers (or equivalently of directed edges in the de Bruijn graph shown in Fig. 3a) in 250 realizations of the system with different values of λ_*ij*_ and *c*_*i*_. Fig 3d shows the distribution of the number (*N* + *N*_*sc*_)/2 of edges in undirected graphs such as one shown in Fig. 3b for these realizations. As discussed above, the deviation of this last quantity down from *Z* is equal to the number of trees in the pseudoforest. As shown in Fig. 3d, it follows approximately exponential distribution with the average around 1.6 (the red line in Fig. 3c). At the same time, the distribution of the number of surviving 2-mers (*N*) has a peak around 36 < 2*Z* = 40. The quantity 2*Z* − *N* is always positive and approximately follows a Poisson distribution with the average of 5.5 (red line in Fig. 3c).

**Figure 3:**
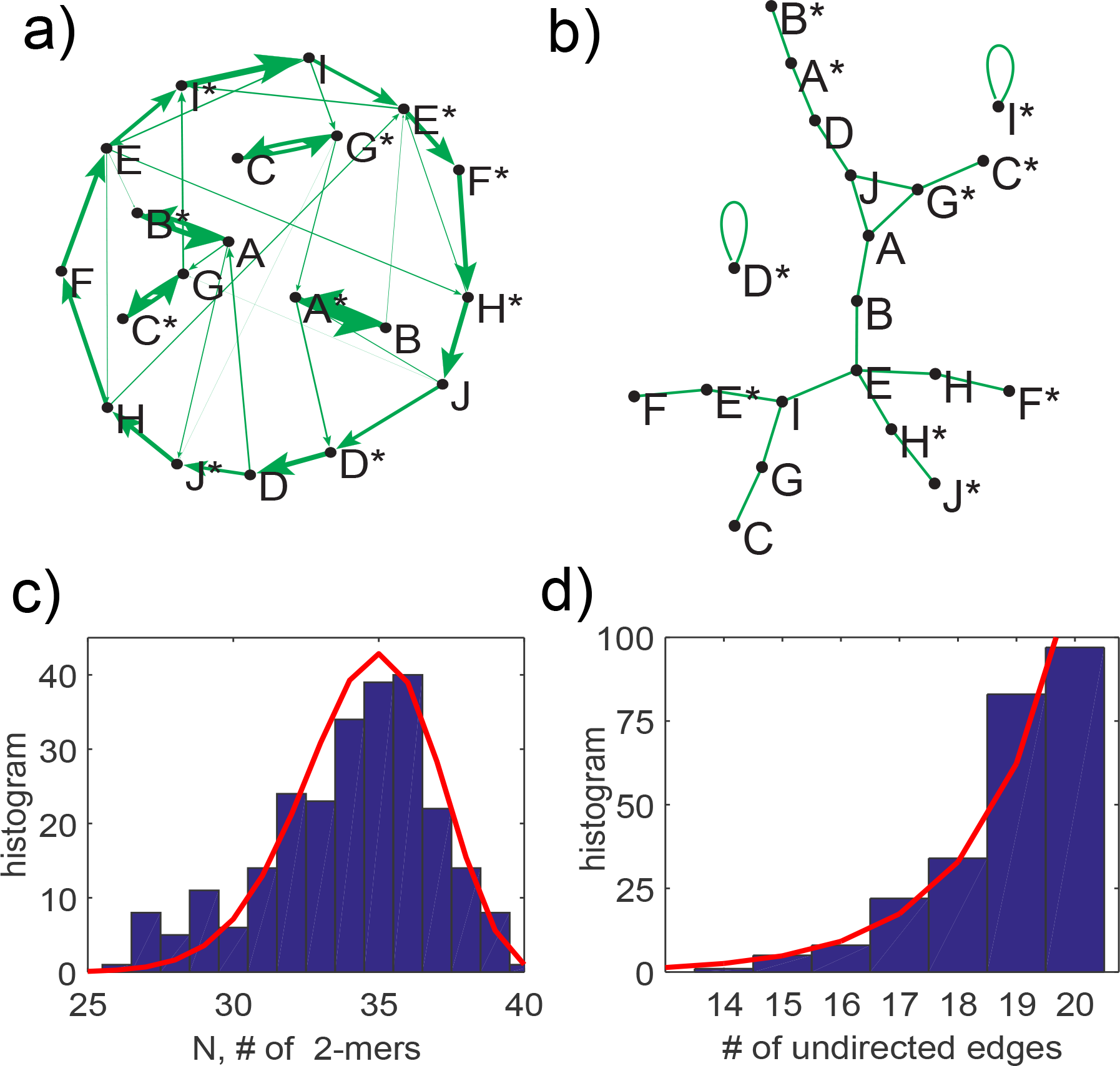
a) The de Bruijn directed graph with *Z* = 20 nodes corresponding to monomers and edges corresponding to surviving 2-mers. Thickness of each edge scales with 2-mer’s concentration. Polymer sequences are walks on this graph. b) Undirected graph representation of the system constructed as described in the text. Each edge represents two mutually complementary 2-mers, for instance, *F* − *E*\* stands for *F* → *E* and *E*\* → *F*\*. c) The histogram of the number *N* of surviving 2-mers (directed edges in the de Bruijn graph). d) The histogram of the number of undirected edges (*N* + *N*_*sc*_)/2 in 250 realizations of the model with different λ’s and *c*_*i*_.

### Variability of the surviving set of 2-mers

The set of surviving 2-mers and their concentrations depend on a number of parameters: ligation rates λ_*ij*_, total monomer concentrations *c*_*i*_, and, possibly, seed concentrations of individual 2-mers.

We analyzed the sensitivity of the steady state of our system with respect to all of these parameters one-by-one. First we fixed both λ_*ij*_, and *c*_*i*_ and analyzed the final 2-mer concentrations for a large number of random realizations of *Z*^2^ small (but positive) seed concentrations. We found the final state to be completely reproducible as long as all seed concentrations are non-zero. Note that due to autocatalytic nature of 2-mer dynamics, Eq. 1, a pair of complementary 2-mers with zero seed concentrations would never emerge on their own.

Next, we fixed the ligation rates λ_*ij*_ to their values used to construct the heatmaps shown in Fig. 2 and networks in Fig. 3. We then simulated 100 realizations of the system with *c*_*i*_ pulled from a log-normal distribution *P*(*c*_*i*_) ~ exp(− [log(*c*_*i*_/*c*)]^2^/*σ*_*c*_)/*c*_*i*_ with *c* = 3 and *σ*_*c*_ = 0.1. Fig. 4a shows the heatmap of the fraction of realizations of *c*_*i*_ in which each individual 2-mer survives in the steady state. In Fig. 4b we present the same results in the form of the histogram (blue bars). The majority of 2-mers (324 out of 400 visible as the leftmost bar in Fig. 4b) got extinct in all of 100 realizations. While the number of surviving 2-mers in each realization never exceeded 2*Z* = 40 (see Fig. 2c), the overall number of 2-mers that survived in at least one realization of *c*_*i*_ was substantially larger: 76. Out of them, 20 “universal survivors” were present in all 100 realizations. Furthermore, as can be seen by comparing heatmaps in Figs 2d and Fig. 4a, these 20 2-mers typically have high steady state concentrations *d*_*ij*_. To further investigate the correlation between 2-mer concentration and its survivability, we analyzed a larger set of 250 realizations of the system with the same fixed ligation matrix but variable monomer concentrations *c*_*i*_. The resulting distribution of 2-mer concentrations shown in Fig. 4c is clearly bimodal. The bimodality is also apparent from an example of de Bruijn graph shown in Fig. 3a, where approximately half of all links (thick lines) correspond to more abundant 2-mers, while the other half (thin lines) - to 2-mers present at low concentrations. The high-concentration peak of the distribution in Fig. 4c is dominated by the contribution from 20 universal survivors shown as red line. Note that despite an increased number of realizations, the set and number of universal survivors stayed the same as in Fig. 4a-b.

We further investigated the variability of the set of surviving 2-mers as a function of width *σ*_*c*_ of the log-normal distribution of monomer concentrations. As shown in Fig. 4d, the number of universal survivors (red line) systematically decreases with *σ*_*c*_, ultimately reaching 0 for *σ*_*c*_ ≥ 0.75. Consistent with this trend, the bimodality of the concentration distribution in Fig. 4c also disappears for larger valu*σ*_*c*_. Note that all numbers of 2-mers shown in Fig. 4d were normalized by 2*Z*, i.e. the upper bound of the number of surviving 2-mers in each realization. The average number of survivors in a single realization (green line) does not have a notable dependence on *σ*_*c*_. The blue line in Fig. 4d shows the number of 2-mers (normalized by 2*Z*) that survived at least once among 100 realizations of *c*_*i*_. Note this curve grows significantly with *σ*_*c*_ reaching the value as high as 5. This corresponds to half of all *Z*^2^ = 400 possible 2-mers having a chance to survive at in least one of these realizations.

**Figure 4:**
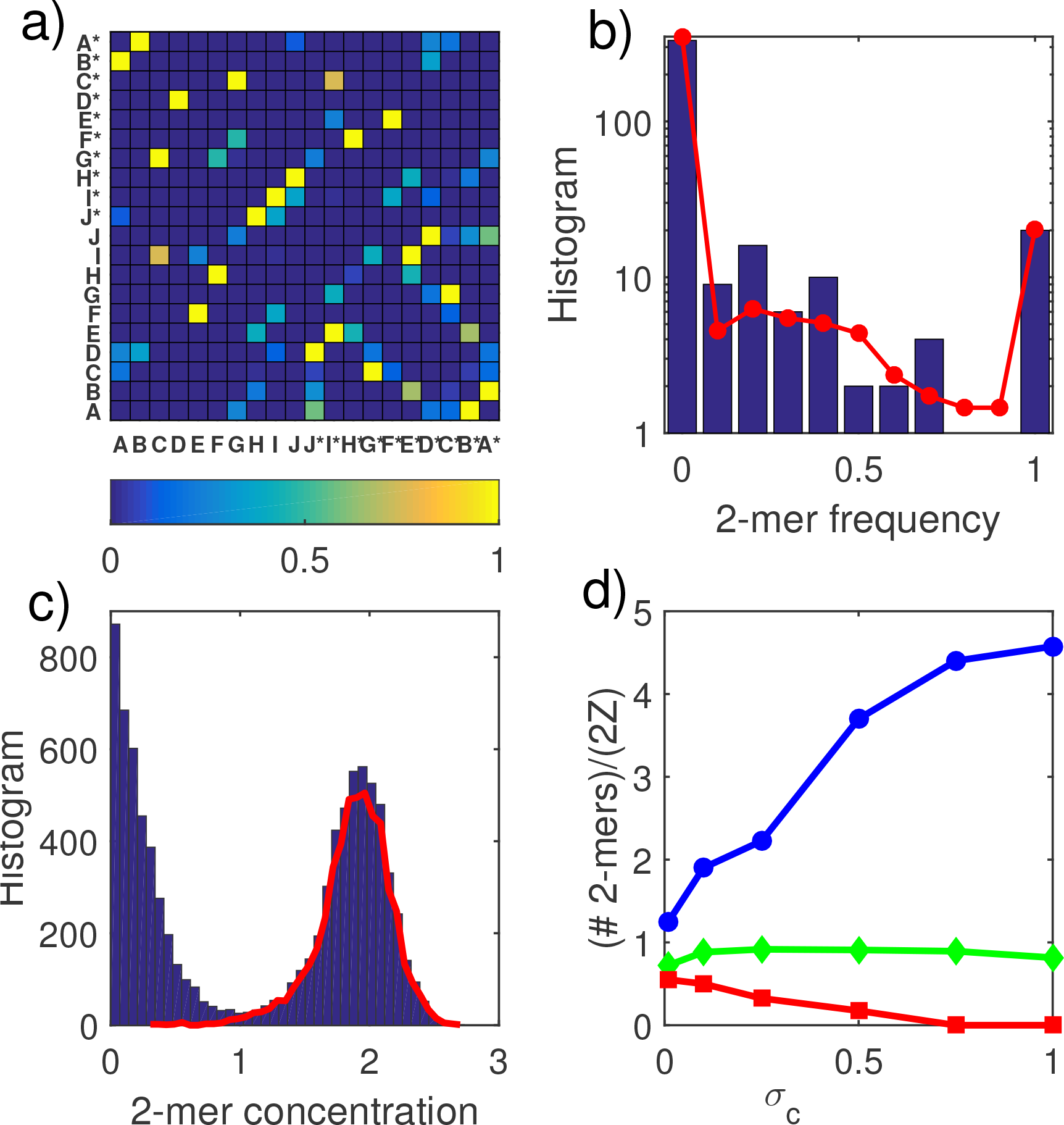
a) The heatmap of survival frequency of 2-mer in the steady state of the system with fixed λ_*ij*_ (same as in Figs. 2-3) and 100 different realizations of *c*_*i*_. Note the similarity to Fig. 2c. b) The histogram of these survival frequencies (blue bars) as well as its average (red line) over 10 different realizations of λ_*ij*_. c) The histogram of 2-mer concentrations in 250 realizations of *c*_*i*_ for fixed λ_*ij*_. The red line shows the contribution from 20 “universal survivors” - 2-mers present across all realizations. d) The number of universal survivors (red line), the average number of survivors in a single realization (green line) and the number of 2-mers present in at least one among hundred realizations of *c*_*i*_ (blue) each normalized by 2*Z*. The x-axis is the width *σ*_*c*_ of log-normal distribution of monomer concentrations.

## Discussion

The major conclusion following from this study is that our model system of mutually catalyzing heteropolymers has a natural tendency towards spontaneous reduction of the information entropy. This represents an effective *reversal* of the second law of thermodynamics in this class of systems. While “*violations*” of the second law are indeed expected in externally driven non-equilibrium systems, the observed “reversal” has much greater implications. Both living organisms and other self-organized systems such as human culture, economics, and technology, are characterized by an ever increasing complexity, indicating the ongoing reduction in the information entropy.

The thermodynamic entropy of a system of heteropolymers is composed of two distinct parts [28] : (i) the translational and configurational entropy of polymer chains; and (ii) the information entropy associated with sequence statistics. Our current model hints at a hierarchical scenario of entropy reduction in populations of heteropolymers. First, the translational entropy is reduced due to template-based polymerization as studied in our previous work [26] within Random Sequence Approximation (RSA). Then RSA breaks down at the level of 2-mers due to their competition with each other for a limited resource of monomers. Such symmetry breaking in the sequence space results in a dramatic reduction in the information entropy (ii). At this point, sequences of the entire pool of chains are generated as Markovian random walks on the de Bruijn graph (Fig. 3a). One can imagine a further reduction of the information entropy due to the emergence of correlations between consecutive steps of this walk.

There are multiple physical scenarios outside the scope of our current model that would lead to such longer-range correlations in the sequence space. They include e.g. a dependence of association rates in Eq. 1 on lengths of the three chains involved in the process of template-assisted ligation. Another intriguing scenario is a spontaneous emergence of chains with weak catalytic activity above and beyond their role as templates for ligation. For instance, some sequences may facilitate breakage or conversely promote ligation reactions, either among some specific sequences or universally. Such sequences would provide a missing link between the prebiotic soup considered here and the emergence of the first ribozymes in the RNA world scenario.

A common pattern in functional RNA-based structures such as ribozymes and ribosomes is the presence of hairpins and loops. Note, however, that mass-action term in the Eqs. 1 assumes that the template and the two substrates belong to three different chains thus ignoring these higher order structures. A proper model description taking them into account is a natural next step in development of our approach. An important future question to address is whether our system would naturally evolve towards or away from loops and hairpins. One might expect that a hairpin composed from several pairs of mutually complementary 2-mers would have a self-healing property: any broken bond would be quickly repaired since the template and one of the substrates belong to the same chain and hence are more likely to be hybridized.

An interesting feature of the sequence statistics emerging in our model is that the entropy is not reduced to its absolute minimum that would correspond to a unique “master sequence”. In the de Bruijn representation such master sequence would look e.g. like a single cycle (or several unconnected cycles) in which for every monomer there is unique right neighbor following it on every chain. In this case the walk on the de Bruijn graph would be completely deterministic. In contrast, in our model each monomer typically has two possible right neighbors in de Bruijn graph. In the limit of relatively small variations *σ*_*c*_ = 0.1 used to construct the Fig. 3a, stronger links corresponding to “universal survivors”, produce *c*_*i*_-independent backbone of the graph akin to the master sequence. Conversely, weaker links allow for infrequent hopping between different parts of the graph leading to deviations from this master sequence. The opposite limit of large variations in monomer concentrations is especially promising from the point of view of further evolution. In that limit, there is no dominant master sequence. This dramatically expands the explored region in the sequence space: now, for every step of the Markovian walk there are on average two comparable probabilities for selecting the next monomer. As a result, the number of possible sequences of length *L* in our model, ~ 2^*L*^, remains exponentially large, yet dramatically reduced compared to its random sequence limit, *Z*^*L*^. Furthermore, in the limit of large *σ*_*c*_, different realizations of *c*_*i*_ give rise to significantly different sets of surviving 2- mers. Note that, unlike the ligation rates λ_*ij*_, monomer concentrations *c*_*i*_ (or equivalently their influxes *ϕ*_*i*_) could vary significantly from one spatial location to another (See [29] for a study of spatial inhomogeneity in the prebiotic context). This allows for an effective exploration of various regions (of size ~ 2^*L*^ each) of the global sequence space, rather than converging to the same subset of sequences over and over.

Our model describes a simple yet general mechanism for spontaneous entropy reduction in systems capable of template-assisted ligation. There are multiple possible experimental realizations of such systems based on either traditional DNA/RNA biochemistry or artificial micro/nano structures. The most direct implementation of our model would be a system composed of *Z* words made of a string of nucleotides bound together by strong (e.g. DNA-type) bonds. They have to be designed to form *Z*/2 mutually complementary pairs that are orthogonal to each other, i.e. words from different pairs have no long overlaps. These words would play the role of composite monomers in our model that could be subsequently connected to each other with weaker, breakable (e.g. RNA-type) bonds. As discussed earlier, our model is directly applicable to the scenario in which all bonds are unbreakable, while the whole system is uniformly diluted and fresh (unbound) monomers are supplied at a constant rate. This greatly expands possibilities for its experimental implementation. Similarly, our dynamics can be implemented e.g. using the DNA origami nanoblocks introduced in Ref. [6]. It should be emphasized, that in order to achieve the behavior described by our model, the experiments need to be conducted well below the saturation regime, i. e. the night phase should be shorter than a typical association time for hybridization.

Yet another interpretation of our model does not involve any long polymers at all. In this case 2-mers are represented by physical dimers made of only two monomeric units incapable of forming longer chains. Our model predicts that, even in this simple system, the compositional entropy would drop because of the extinction of most of the dimers leaving only *N* ≤ 2*Z* survivors. On the one hand, this further extends possibilities for experimental implementation. For instance, one could construct the DNA-based system described above but limited to no more than 2-word chains. This would greatly reduce the complexity of the screening process.

On the other hand, this dimer interpretation has an intriguing connection to the Kauffman model of autocatalytic chemical reaction networks [12]. In the Kauffman case, some of the molecules types in the pool are capable of catalyzing the synthesis of others from “raw materials” (abundant small metabolites) ultimately resulting in the emergence of metabolic auto-catalytic cycles in large systems. A recent model of such chemical reactions [30] shows that the system self-organizes to a state finely tuned to the external driving force. This can be interpreted as maximization of the rate of negentropy adsorption from the environment. In our case dimers correspond to mutually catalytic entities while monomers represent raw materials. While in the current implementation of our model, the catalytic cycles could only involve two mutually complementary dimers, it is straightforward to generalize the model to allow the mutual catalysis of any pair of dimers. We expect our findings about the reduction of entropy to be fully transferable to that case. Thus, our model has a potential of bridging the gap between two traditionally competitive visions of the Origin of Life: “information first” and “metabolism first”.

## Acknowledgements

This research used resources of the Center for Functional Nanomaterials, which is a U.S. DOE Office of Science User Facility, at Brookhaven National Laboratory under Contract No. DE-SC0012704.

